# Detection and treatment strategy for *Tritrichomonas muris* in the common laboratory mouse

**DOI:** 10.1101/827055

**Authors:** Andreia S. Da Costa, Tisha M. Graham, Jennifer A. Duncan, Smitha P.S. Pillai, Jennifer M. Lund

## Abstract

Maintaining a specific pathogen-free (SPF) mouse colony is critical to avoid potentially confounding variables introduced by unknown infections. Here, we report an instance of protozoan *Tritrichomonas muris* (*T. muris*) infection of mice within an SPF facility. Although *T. muris* has been considered a commensal organism, we observed instances of asymmetric infection in gene-knockout mice as compared to wild-type mice. We utilized treatment with metronidazole and confirmed successful elimination of *T. muris* from our SPF colony using extraction of fecal DNA followed by PCR detection. We propose that T. muris testing should be considered for SPF mice, particularly in immunity studies.

## Introduction

*Tritrichomonas muris* is a triflagellate single-celled protozoan that infects the intestinal tissues of mice and other rodents. It is closely related to *Tritrichomonas foetus*, a pathogen known to infect bovine reproductive organs (Yao and Koster, 2015). *T. muris*, however, has been listed as a commensal organism (Baker, 2008) and is therefore rarely screened for during routine mouse colony maintenance. Nevertheless, a proteomic analysis of intestinal tissue collected from mice infected with *T. muri*s differentially expressed several proteins compared to uninfected intestines, a subset of which were immune related (Kashiwagi et al., 2009), suggesting that *T. muris* infections may trigger an immune response. More recently, a study of T-cell-driven colitis identified that *T. muris* infection exacerbates disease and skews baseline T-cell homeostasis toward a more pro-inflammatory mucosal environment (Escalante et al., 2016), and a second study also found that host-protozoan interactions can alter mucosal immune homeostasis (Chudnovskiy et al., 2016). *T. muris* infection has been shown to increase the frequency of intestinal tuft cells which can induce IL-13 secretion from intestinal innate lymphoid cells (ILC) (Howitt et al., 2016; von Moltke et al., 2016). Additional studies showed that a metabolite from *T. muris* trigger tuft cell production of IL-25 (Schneider et al., 2018).

During a routine intestinal tissue preparation, we discovered that a subset of our mouse colony was infected with *T. muris.* Specifically, we determined that a conditional knock-out (cKO) mouse strain was most affected, and so it was critical that we eliminate the potential confounding effects of a *T. muris* infection in our cKOs for accurate comparison to WT controls. Thus, we developed a PCR-based detection assay to determine the *T. muris* infection status of our cKO mouse model in conjunction with a metronidazole-based treatment plan that allowed us to re-establish the SPF status of our colony.

## Results

### T. muris infection can potentially confound characterization of novel immune system-associated gene deletion mouse models

Gene knockout (KO) and Cre/*lox* conditional knockout (cKO) mouse models have proven to be invaluable tools in the reductive investigation of individual gene functions in specific cells and tissues. Use of KOs in studies of the immune system vary widely and have helped elucidate the inner workings of immune cells as well as their relationships with other systems within the organism (Mak et al., 2001). Accordingly, we set out to characterize the effect of integrin β1 deletion in regulatory T cells (Treg) using Itgb1^*flox*/flox^ x Foxp3^*Cre*^ (cKO) mice (Da Costa et al., 2019). When preparing small intestine from cKO and WT control mice for the analysis of lamina propria lymphocytes, we observed a dramatic discoloration in the cKO-derived tissue (Fig. 1A) and subsequently detected a free-swimming organism in the tissue supernatant under 20X magnification. Fecal samples from cKO mice were sent for identification of the protozoan (IDEXX Bioresearch, Columbia, MO, USA), which was confirmed to be *Tritrichomonas muris*. Although *T. muris* has been grouped with other protozoan infections associated with symptoms including weight loss, runting and diarrhea (Charles River Laboratories International, 2009), we did not find a statistically significant weight difference in cKO mice as compared to age-matched WT controls (Fig. 1B). To confirm infection, we performed histopathological evaluation of sections of the gastrointestinal tract. Within the lumen of cecum and colon in the cKO group, there were large numbers of oval to pyriform protozoal trophozoites, measuring approximately 20×10μm, with eosinophilic cytoplasm and round hyperchromatic nuclei consistent with *Trichomonas* species (Fig. 1C, left panel). No changes in mucosal inflammation were detected. In the control mouse (Fig. 1C, right panel), no protozoa were detected. Spleen sections were also evaluated from both groups, though we found no appreciable histological differences between *T. muris*-infected cKO mice and uninfected WT controls (Fig. 1D). Altogether, we found evidence of select *T. muris* infection of the intestine of a cKO mouse strain compared to WT mice within an SPF mouse colony.

**Figure 1.**
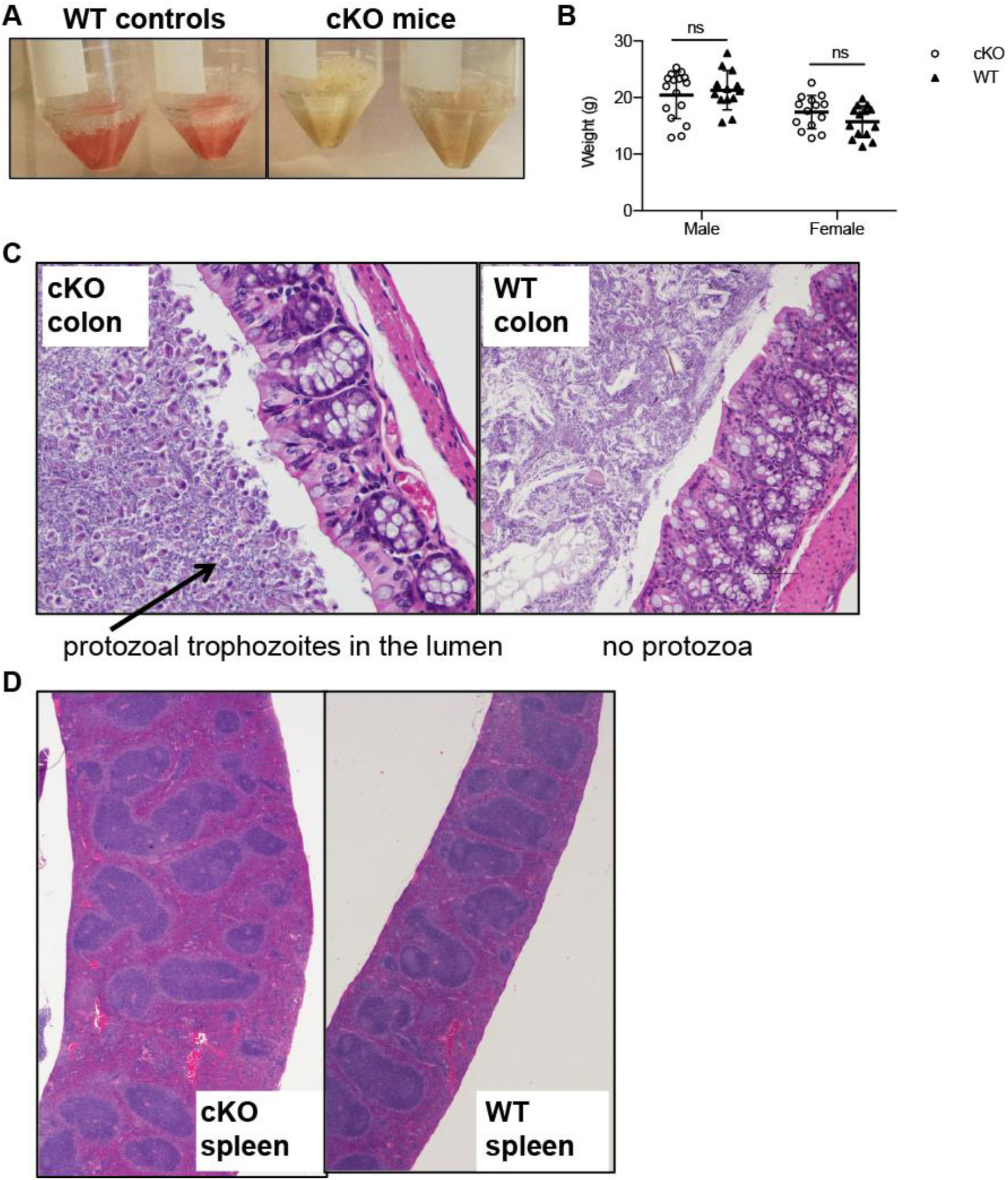
T. muris infection can confound characterization of novel immune system-associated gene deletion mouse models. (A) Left panel shows excised and sectioned small intestine from two adult C57BL/6J male mice in 1X RPMI media supplemented with 5% heat inactivated fetal bovine serum (HI-FBS) after incubation at 37°C with agitation at 220 rpm for 30 mins in EDTA Buffer. Original image contrast has been increased by 40% to improve visibility. Right panel shows small intestine tissue after processing as in left panel, however tubes contain tissue from adult cKO that are infected with *T. muris.* (B) Weights were recorded from male and female cKO and WT on C57BL/6J background. Average weight/age of cKO (n=16) males: 20.4g/52.6 dys, cKO (n=14) females: 17.4 g/48 dys; WT (n=13) males: 21.3g/53.2 dys, WT (n=14) females: 15.7g/48.4 dys. Plots display range and median for each group. Statistics represent two-stage linear step-up procedure of Benjamini, Krieger and Yekutieli, with Q = 1%. Comparison groups (cKO vs. WT) were analyzed individually, without assuming a consistent SD. (C) Histopathological evaluation of colon from cKO or WT mice. Arrow in left panel indicates protozoal trophozoites which are absent in colon tissue assessed from WT mice in the right panel. (D) Histopathological evaluation of spleens from cKO and WT mice. No gross or histological differences were noted between infected cKO and uninfected WT mice.

### PCR-based detection of T. muris reveals wide-spread but not uniform infection of SPF mice

In order to develop the most effective PCR primers given the lack of publicly available sequence data for *T. muris*, we aligned the 18s ribosomal sequences of *T. muris* and *T. foetus* using the nucleotide blast program hosted by NCBI BLAST, and selected primers that covered conserved exon regions shared between both species. Using these primers for PCR, we sought to determine the extent of *T. muris* infection within our cKO breeder population and colony. To ensure that infection status was attributed correctly to individual mice, each mouse tested was segregated to a clean receptacle until a fresh fecal sample was produced. We found that each cKO breeder cage sampled tested positive for *T. muris* DNA (Fig. 2A lanes 3-10). Furthermore, Itgb1^*flox*/wt^ x Foxp3^*Cre*^ mice also tested positive (Fig. 2A, lanes 1 and 2). This indiscriminate infection pattern suggests that the gene deletion in the cKO mice did not result in an immune deficit that made them uniquely susceptible to *T. muris*. Another cKO strain, CTLA4^*flox/flox*^ x Foxp3^*Cre*^, was previously tested commercially and so was utilized successfully as a negative infection-control (Fig. 2A, lanes 11 and 12). After determining complete infection of Itgb1^*flox/flox*^ x Foxp3^*Cre*^ breeding stock, we further confirmed that the infection had spread to other strains despite the lack of physical contact (Fig. 2A, lanes 14-20). Surprisingly, the sentinel mice housed in the cKO rack tested negative for *T. muris* DNA (Fig. 2A, lane 21). As sentinel mice are routinely exposed to bedding from various cages housed in the same rack, therefore the lack of *T. muris*, infection in these mice underscores the need to expose sentinels to fresh feces to most effectively surveil a colony.

**Figure 2.**
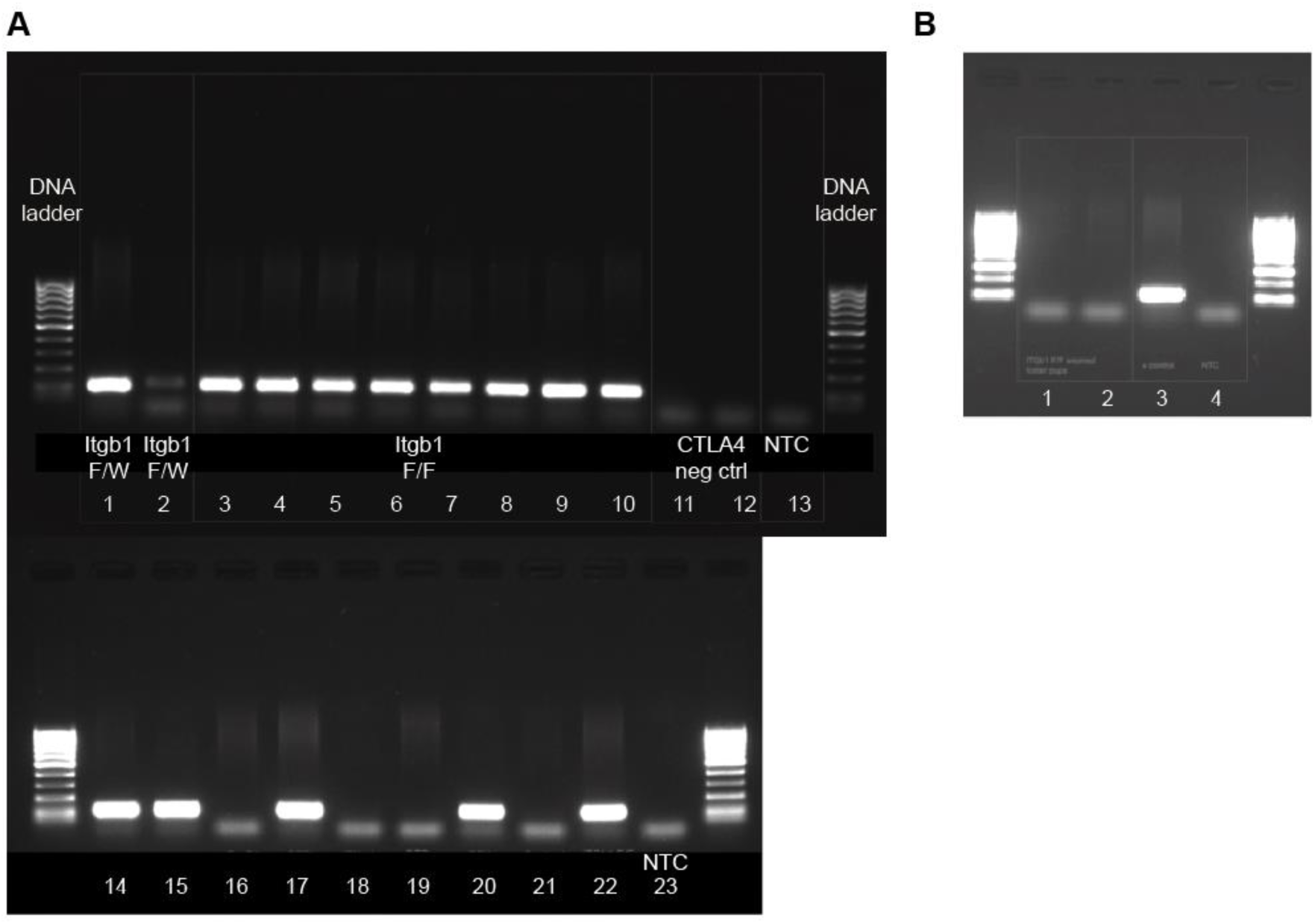
T. muris specific primer selection and PCR-based detection of T. muris infection. (A) Using primers and PCR protocol described, we collected fecal samples from all breeder cages housing our Itgb1^F/F^ x Foxp3^Cre^ and Itgb1^F/W^ x Foxp3^Cre^ control mice and found that all breeder cages were positive for presence of *T. muris* DNA (lanes 1-10). CTLA4 (Ctla4^F/F^ x Foxp3^Cre^) are a mouse strain known to be negative through commercial testing (IDEXX Bioresearch, Inc) (lanes 11-12). NTC = non-template control where template has been replaced with additional DNase-free dH_2_0 (lane 13). Additional strains within our colony were tested and some were found to be infected (lanes 14-20). Long-lived sentinel mice associated with the contaminated rack tested negative (lane 21). (B) cKO foster pup fecal samples tested negative for *T. muris* DNA after metronidazole treatment (lanes 1-2). + control = DNA extracted from fecal samples of CKO mice prior to treatment (lane 3). NTC = No template control (lane 4).

### Metronidazole treatment in combination with a fostering plan can clear T. muris infection

To avoid the costly and time-consuming, albeit guaranteed strategy of replacement of all affected cages by purchase of new breeder stock, we used a combination of metronidazole treatment in combination with a pup-fostering plan. Metronidazole serves as both an antibiotic and antiprotozoal medication often prescribed to treat human *Giardia lamblia*, *Clostridium difficile* and *Trichomonas vaginalis* infections (Freeman et al., 1997). Although very little has been published describing treatment strategies for *T. muris*, Roach *et al* showed that *T. muris* introduced from infected gerbils to mouse hosts was successfully treated by supplementing drinking water with metronidazole or tinidazole (Roach et al., 1988).

Thus, we purchased outbred Swiss Cr1:CD1 (ICR) breeders both for ease of differentiation of foster pups by coat color, and due to anecdotal reports of CD-1 dams’ efficient handling of large litter sizes. CD-1 mice were tested upon arrival and found to be negative for fecal presence of *T. muris* DNA. CD-1 and cKO breeder pairs were subsequently set-up on different racks. After visual confirmation of the development of a mucus plug in pregnant females, breeder males were removed from the breeding cages. Gestating dams were then monitored closely and metronidazole-supplemented water introduced approximately 5 days prior to expected birth. Upon birth, CD-1 pups were removed from the breeder cage and replaced with the recently birthed cKO pups. Metronidazole supplementation was continued for 5 days post-pup transfer. After a total of 10 days of water supplementation, metronidazole treatment was ceased and pups remained under the care of the CD-1 foster dam until 21 days post-birth. Foster pups were weaned, fecal samples collected and DNA extracted. PCR analyses of the collected fecal samples were negative for the presence of *T. muris* DNA, indicating successful treatment (Fig. 2B). Although several studies, both mouse and human, have concluded that metronidazole does not increase the incidence of cancer in treated patients (or animals) (Chacko and Bhide, 1986; Thapa et al., 1998), the Food and Drug Administration (FDA) has labeled metronidazole as a potential mutagen and teratogen. Thus, we completed several rounds of backcrossing of the newly fostered *T. muris*-free cKO mice to WT C57BL/6J mice, both to limit the carry-over of potential mutagenic effects of metronidazole treatment and to mitigate the effects of genetic bottlenecking as a result of the necessarily small number of pups fostered. Taken, together, we report a strategy to test and treat for *T. muris* infection in mice.

## Discussion

We describe a treatment strategy developed in response to the incidental finding of *T. muris* infection within select strains in our mouse colony. To ensure the accurate and high-throughput identification of the protozoa, we developed a PCR-based assay to evaluate the presence of *T. muris* DNA in fresh fecal samples. Through this testing, we determined that the infection had spread through the colony and that the use of sentinel mice was not sufficient to detect *T. muris* infection. Due to our finding of differential *T. muris* infection of the intestine between cKO and WT mice (Figs. 1C and 2A) in which we wished to analyze immune cell phenotypes, along with the knowledge that *Tritrichomonas* infections have a demonstrated effect of altering mucosal immune homeostasis (Escalante et al., 2016; Howitt et al., 2016; Schneider et al., 2018; von Moltke et al., 2016) that could thus confound our studies, we elected to eradicate the infection from our colony. We developed a treatment plan that resulted in the successful restoration of the original controlled SPF-status of our cKO mice, thereby eliminating a potential driver of differential immune homeostasis between mouse strains.

The regulated exposure of experimental lab animals to a pre-determined list of pathogens and commensals has become a hallmark of rigorously conducted studies. Ensuring the reliability and integrity of data generation within animal models requires a thorough understanding of potential confounding variables. This is especially important in studies that propose to investigate the complex subtleties of a developing immune response. Furthermore, studies that include the characterization of a novel gene deletion in a mouse model are particularly vulnerable to misdirection by unknown confounding variables. As such, it is critical that investigators remain vigilant of the microbial status of their colonies as well as remaining open to the re-evaluation of micro-organisms that may have been deemed commensals in the past. Based on our study, we propose that *T. muris* should be monitored as a pathosymbiont in SPF colonies, particularly if mice are to be used for the study of immunity, since infection of a KO but not a control could lead to a mis-classification or mis-interpretation of any differential data.

## Methods

### Ethics Statement

All animal work was approved by the Fred Hutchinson Cancer Research Center (Fred Hutch) and treatment provided through the Comparative Medicine department. The Office of Laboratory Animal Welfare of the National Institutes of Health has approved Fred Hutch (#A3226-01), and this study was carried out in strict accordance with the Public Health Service Policy on Humane Care and Use of Laboratory Animals.

### T. muris specific PCR amplification and visualization

Fresh fecal pellets were collected from mice and fecal DNA extracted using the ZR Fecal DNA MiniPrep™ kit (Zymo Research Corp., CA, U.S.A.). *T. muris* forward and reverse primers were purchased through Integrated DNA Technologies (IDTDNA.com). Primer sequences are as follows: forward primer, GCAATGGATGTCTTGGCTTC, and reverse primer, GCGCAATATGCATTCAAAGA. Master mix composition was based on manufacturer’s recommendations included in the 2X MyTaq™ HS Red Mix purchased through Bioline USA Inc, (Taunton, MA). Annealing temperature, duration and cycle number were determined empirically: Tm (50mM NaCl) for forward primer = 54.4°C; Tm (50mM NaCl) for reverse primer = 51.9°C. Gel electrophoreses of PCR products were done using 1% Agarose gel matrix with the addition of 10,000X GelRed™ Nucleic Acid Gel Stain (Biotium, Inc. CA, USA). Samples were run alongside 100bp exACTGene™ DNA ladder (Fisher BioReagents™) for ~35mins at 100V in 1X TAE Buffer. Expected PCR product is approximately 65bp in length.

### Metronidazole treatment

An optimized dose of 2.5 mg/ml metronidazole in a 1% sucrose solution was used to treat *T. muris* in mice(Roach et al., 1988). A solution of 2.5 mg/ml metronidazole was prepared by crushing a 250mg metronidazole tablet and dissolving it in 100ml of a 1% w/v sucrose solution. The solution was then passed through a 22um filter and placed in sterile water bottles wrapped in foil to prevent photo-degradation.

### Mice

C57BL/6J;129-Itgb1^Flox/Flox^ x Foxp3^eGFP-Cre-ERT2^ mice were a generous donation from Dr. Alexander Rudensky (Memorial Sloan Kettering Cancer Center) and have been maintained at the Fred Hutch Animal Facility for 10+ years. Six-week old Crl:CD1 (ICR) foster dams were purchased from Charles River.

### Histopathology

Complete necropsies were performed on wild-type (WT) and cKO mice. Tissues including spleen, liver, and intestines were collected and fixed in 10% neutral buffered formalin, routinely processed and embedded in paraffin, sectioned at 4µm, and stained with hematoxylin and eosin for histological evaluation.

## Acknowledgements

The authors thank Rajesh Uthamanthil, DVM, MVSc, PhD, DACLAM for advice and the Fred Hutchinson Cancer Research Center Dept. of Comparative Medicine for assistance and guidance.

## Author Contributions

Development and execution of treatment strategy (A.S.D., T.M.G, J.A.D.); Histopathology scoring, interpretation and analysis (S.P.S.P); Writing and preparation of manuscript (A.S.D. and J.M.L.); Critical revisions and guidance (J.M.L).

## References

Baker, D.G. (2008). Parasites of rats and mice, in Flynn’s Parasites of Laboratory Animals. Blackwell Publishing Ltd Second Edition, 303–397.

Chacko, M., and Bhide, S.V. (1986). Carcinogenicity, perinatal carcinogenicity and teratogenicity of low dose metronidazole (MNZ) in Swiss mice. J Cancer Res Clin Oncol 112, 135–140.

Charles River Laboratories International, I. (2009). Intestinal Protozoa in Rodents: Technical Sheet. Charles River Research Models and Services, 1–2.

Chudnovskiy, A., Mortha, A., Kana, V., Kennard, A., Ramirez, J.D., Rahman, A., Remark, R., Mogno, I., Ng, R., Gnjatic, S., et al. (2016). Host-Protozoan Interactions Protect from Mucosal Infections through Activation of the Inflammasome. Cell 167, 444–456 e414.

Da Costa, A.S., Graham, J.B., Swarts, J.L., and Lund, J.M. (2019). Regulatory T cells limit unconventional memory to preserve the capacity to mount protective CD8 memory responses to pathogens. Proc Natl Acad Sci U S A.

Escalante, N.K., Lemire, P., Cruz Tleugabulova, M., Prescott, D., Mortha, A., Streutker, C.J., Girardin, S.E., Philpott, D.J., and Mallevaey, T. (2016). The common mouse protozoa Tritrichomonas muris alters mucosal T cell homeostasis and colitis susceptibility. J Exp Med 213, 2841–2850.

Freeman, C.D., Klutman, N.E., and Lamp, K.C. (1997). Metronidazole. A therapeutic review and update. Drugs 54, 679–708.

Howitt, M.R., Lavoie, S., Michaud, M., Blum, A.M., Tran, S.V., Weinstock, J.V., Gallini, C.A., Redding, K., Margolskee, R.F., Osborne, L.C., et al. (2016). Tuft cells, taste-chemosensory cells, orchestrate parasite type 2 immunity in the gut. Science 351, 1329–1333.

Kashiwagi, A., Kurosaki, H., Luo, H., Yamamoto, H., Oshimura, M., and Shibahara, T. (2009). Effects of Tritrichomonas muris on the mouse intestine: a proteomic analysis. Exp Anim 58, 537–542.

Mak, T.W., Penninger, J.M., and Ohashi, P.S. (2001). Knockout mice: a paradigm shift in modern immunology. Nat Rev Immunol 1, 11–19.

Roach, P.D., Wallis, P.M., and Olson, M.E. (1988). The use of metronidazole, tinidazole and dimetridazole in eliminating trichomonads from laboratory mice. Lab Anim 22, 361–364.

Schneider, C., O’Leary, C.E., von Moltke, J., Liang, H.E., Ang, Q.Y., Turnbaugh, P.J., Radhakrishnan, S., Pellizzon, M., Ma, A., and Locksley, R.M. (2018). A Metabolite-Triggered Tuft Cell-ILC2 Circuit Drives Small Intestinal Remodeling. Cell 174, 271–284 e214.

Thapa, P.B., Whitlock, J.A., Brockman Worrell, K.G., Gideon, P., Mitchel, E.F., Jr., Roberson, P., Pais, R., and Ray, W.A. (1998). Prenatal exposure to metronidazole and risk of childhood cancer: a retrospective cohort study of children younger than 5 years. Cancer 83, 1461–1468.

von Moltke, J., Ji, M., Liang, H.E., and Locksley, R.M. (2016). Tuft-cell-derived IL-25 regulates an intestinal ILC2-epithelial response circuit. Nature 529, 221–225.

Yao, C., and Koster, L.S. (2015). Tritrichomonas foetus infection, a cause of chronic diarrhea in the domestic cat. Vet Res 46, 35.

